# Comparative Approaches for Quantification of Product Yield in the Production of Recombinant Green Fluorescent Protein (GFPuv) Expressed in *E. coli*

**DOI:** 10.1101/2024.06.24.600411

**Authors:** Gabrielle Rusch, Junhyeong Wang, Keith Breau, Katie Kilgour, Gary Gilleskie, Jeff Keele, Kurt Selle, Scott T. Magness, Stefano Menegatti, Michael Daniele

## Abstract

Process Analytical Technologies (PAT) are critical for efficient and automated bioprocessing; moreover, sensor and assay-driven automation will be necessary for the realization of Industry 4.0. Herein, we demonstrate methods for analyzing product yield from a pilot-scale bioreactor (300L) and conduct cross-method comparisons. We used a system of recombinant green fluorescent protein (GFPuv) expressed in *Escherichia coli* (*E. coli*), which is a simple, cost-effective model for evaluation at both laboratory and pilot scales. By comparing inline bioreactor measurements, plate reader assays, and a novel image analysis pipeline, we identify optimal harvest timelines and demonstrate the strengths and limitations of each technique. Results indicate peak cell density and GFPuv expression between 12.5h and 18h post-inoculation, with declining viability thereafter. Comparisons across techniques suggest that imaging methods may be more effective in capturing adverse outcomes, such as increased membrane permeability or cell death, which typically occur at later stages of operation. This integrated analysis offers actionable insights for optimizing biomanufacturing workflows and advances the development of scalable PAT applications, bridging the gap between laboratory research and industrial implementation.

## 1. INTRODUCTION

As bioprocessing technologies advance and continue to scale from laboratory settings to industrial operations, the need for robust analytical methods to monitor and control yield becomes increasingly critical. Inline, at-line, and online Process Analytical Technologies (PAT) have emerged as essential tools for real-time monitoring, enabling enhanced process understanding and control [1]. However, the diversity of available PAT methods—spanning spectroscopic, chromatographic, and imaging-based techniques—presents a challenge in determining the most effective approaches for specific applications. Comparative assessments of these technologies are vital to identify their strengths, limitations, and suitability for diverse bioprocessing scenarios, particularly in the context of industrial demands for precision and scalability. Such comparative studies are not only fundamental for optimizing current biomanufacturing workflows but also for laying the groundwork for transformative innovations in Industry 4.0, including digital twins and predictive modeling [2-5].

For example, digital twins, as virtual representations of physical processes, are fundamentally dependent on the continuous flow of accurate and dynamic data, which in turn requires the application of precise and reliable PAT. In order to effectively inform and drive these digital twin systems, it is imperative to develop a robust understanding of how various PAT techniques perform under diverse operational conditions. Such knowledge enables the thoughtful weighting and integration of their outputs into digital twin frameworks and predictive models, ensuring their relevance and efficacy in optimizing biomanufacturing processes. For instance, as the industry progresses toward more sophisticated digital twin systems and predictive modeling, imaging datasets—such as those proposed in this study—will increasingly play a pivotal role, offering critical insights that bridge the gap between real-time process monitoring and advanced computational modeling [6-9].

The use of green fluorescent protein-expressing *Escherichia coli* (GFPuv-*E*.*coli*) represents a versatile and powerful model system for studying bioprocesses and recombinant protein biomanufacturing [10-17]. *E. coli* is widely utilized in the commercial production of a diverse array of proteins, including industrial enzymes, diagnostic reagents, and therapeutic molecules such as insulin, growth factors, and interferons [18, 19]. The inherent scalability and ease of genetic manipulation of *E. coli* solidify its status as a preferred platform for high-yield expression systems, facilitating the efficient biomanufacturing of complex recombinant proteins for both pharmaceutical and biotechnological applications [20]. Induction of GFPuv expression in *E. coli* can be realized by the introduction of isopropyl β-D-1-thiogalactopyranoside (IPTG) or temperature change, which accelerates fluorescent marker expression and proliferation simultaneously; nevertheless, while IPTG and temperature change are popular induction methods, it should be noted that induction conditions are highly dependent on expression system [23-25]. This dual-functionality makes GFPuv-*E*.*coli* an exemplary model for optimizing various biomanufacturing parameters, spanning from induction strategies and nutrient optimization to evaluating the influence of environmental conditions on yield [26-28]. In total, GFPuv-*E*.*coli* offers a close approximation to key aspects of industrial recombinant protein production, such as transcriptional and translational efficiency, metabolic burden, and stress responses. By genetically fusing GFP or GFPuv to the protein of interest or co-expressing it in parallel, researchers can dynamically track both the temporal and spatial aspects of protein production throughout the bioprocess continuum. In the GFPuv-*E. coli* model, GFPuv provides a direct, quantitative measure of protein yield, facilitating rapid assessment of culture conditions. As a highly quantifiable reporter protein, GFP enables real-time monitoring of expression kinetics, cellular proliferation, and metabolic activity, offering critical insights into bioprocess optimization and control. [21, 22, 29-31].

Although GFP is easily measured and GFPuv-*E*.*coli* is widely acknowledged as a reliable model for assessing bioprocess performance, the differences in sensitivity, resolution, and scalability across various analytical techniques for quantifying product yield can lead to discrepancies in the measurement of protein expression levels, complicating efforts to optimize bioprocess parameters. Spectrophotometric techniques, such as inline bioreactor probes, plate reader assays, and microscopy-based measurements, each offer distinct advantages and limitations. Inline probes allow for real-time monitoring but may lack the sensitivity to capture population or spatial heterogeneity [15, 32]. Plate readers facilitate higher-throughput analysis, yet are limited to bulk measurements, potentially missing localized variations within the bioreactor. On the other hand, advanced microscopy techniques offer single-cell resolution and can capture detailed spatial distributions, though scalability for continuous monitoring in industrial processes remains a concern [33]. These differences in methodology can lead to discrepancies in GFP expression levels, underscoring the need to carefully select analytical techniques based on the specific goals and scale of the bioprocess. Addressing these variations is crucial for ensuring reliable and accurate yield assessments, thereby optimizing both research and industrial applications. Combining and comparing these two perspectives of single cell and batch level analysis is important for accurate analysis of the product, while traditional methods of PAT typically focus on one method of analysis [34-38].

This study systematically compares and contrasts multiple approaches for quantifying cell proliferation and GFPuv expression over time, as illustrated in **Figure 1**. Using GFPuv-*E*.*coli* cultivated under industrial conditions, we assessed commonly employed analytical techniques in biomanufacturing. While GFP and these PAT techniques are well-documented in research, their continued evaluation and comparison hold utility for enhancing bioprocessing performance, but more notably for identifying the degree of variation and limitations, which are critical yet underreported needs for realizing and implementing data driven control and prediction innovations, such as digital twins and continuous production paradigms. By comparing and refining quantification methods for GFP yield, this work contributes to the development of more precise, data-driven biomanufacturing systems, supporting their readiness for the future of industrial biotechnology and aligning with the evolving goals of Industry 4.0.

**Fig 1.**
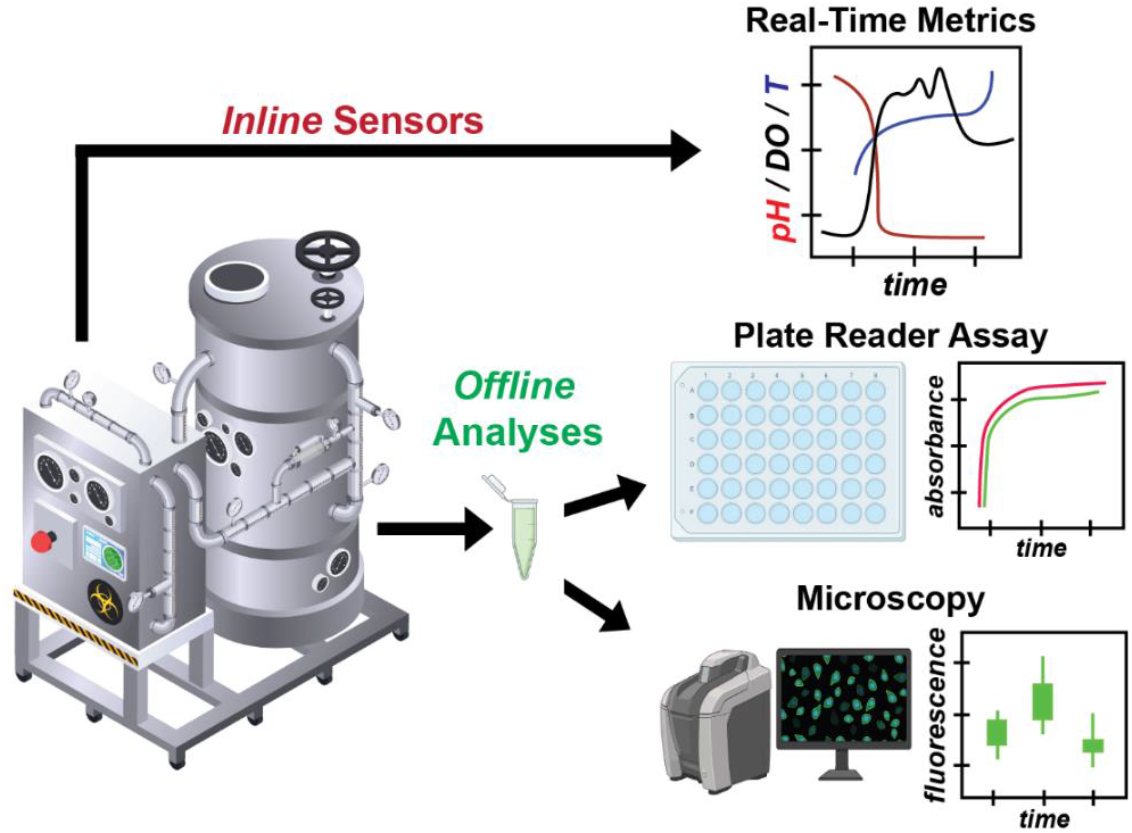
Overview of inline and offline analysis for this bioprocess. Data was collected from several in line sensor modules to be compared against offline analytics including OD600 measurements from a plate reader and confocal image analysis to better understand the bioreactor performance.

## 2. MATERIALS & METHODS

### 2.1 Production of GFPuv-*E.coli* and Sample Collection

A 300 L fed-batch bioreactor used for all processes. Following basic sterilization and preparatory steps, at 0 h, the bioreactor was inoculated with GFPuv-expressing *E*.*coli* (strain = BL21 (DE3)::pET17b::gfpuv). At 6 h, GFPuv expression was induced by introducing isopropyl β-D-1-thiogalactopyranoside (IPTG) into the culture medium. Culture continued for 24h post inoculation and was terminated upon harvest. Inline monitoring was performed throughout the 24-hour bioreactor run to measure several critical parameters, including optical density (OD600), glucose concentration, pH, percentage of dissolved oxygen (DO%), temperature, and system weight. The culture was continuously agitated, with mixing speed (rotations per minute, rpm) and air sparging rate (standard liters per minute, slpm) recorded to ensure consistent aeration and homogeneity within the reactor. Two separate bioreactor runs were conducted. For both bioreactor runs, 2 x 1 mL samples were collected every hour between inoculation and induction. Post-induction, 2 x 1 mL samples were collected every 30 min. Samples were stored at −80°C prior to analysis. The timeline for sample collection is presented in **Figure 3D**.

### 2.2 Plate Reader Assays

OD_600_ and fluorescence intensity were measured on a Cytiva XYB98 plate reader. Fluorescence intensity was measured at λ_ex_ = 395 nm and λ_em_ = 508 nm. Each sample underwent thawing and 100 μL of each sample was dispensed into a black 96-well plate with a clear bottom for analysis. Technical triplicates were prepared and analyzed for each sample from both bioreactor runs. Following sample preparation and initial OD_600_ and fluorescence measurements were recorded, the plates were centrifuged at 4000 xg for 3 min. Post-centrifugation, the supernatant was aspirated and plated alongside the pellet. OD_600_ and fluorescence measurements were then recorded.

### 2.3 Confocal Microscopy and Image Analysis

GFPuv-*E*.*coli* cells were also analyzed using confocal microscopy as a secondary offline analytical technique. Laser scanning confocal microscope (Leica Stellaris8, water immersion 63x) was used for all imaging. Prior to imaging, samples were thawed and 500 μL of each sample was removed, stained with CellBrite 640, fixed using 4% paraformaldehyde (PFA), and washed with phosphate buffered saline (PBS) following standard protocols. After fixing, staining, and washing, each sample was resuspended in 5 mL of PBS and further diluted 1:200 for optimal imaging density. However, sample S1 at the 0 h collection contained a low cell count for both runs, necessitating a 1:20 dilution instead of the standard 1:200 dilution. For ease of imaging multiple samples, basic flow cells were fabricated using polydimethylsiloxane (PDMS) sealed to glass slides as shown in **Figure 2**. Each well of the fabricated flow cell was filled with 25 μL of the prepared and diluted sample. GFP images were collected at λ_ex_ = 488 nm; λ_em_ = 507 nm set at 50.3% gain and 1.5% intensity, while the CellBrite 640 nm membrane stain images were collected at λ_ex_ = 642 nm; λ_em_ = 662 nm adjusted between 16-25% gain and 1.5% intensity for image acquisition. The membrane stain facilitated the identification of individual *E. coli* cells regardless of GFP expression to construct an imaging mask. When collecting CellBrite stain images, the gain and intensity were modified slightly from sample to sample, however the GFP images were collected at the same gain and intensity to allow for appropriate comparison of relative fluorescence per GFPuv-*E*.*coli*. The zoom was set to 1.0 and the pinhole was set to 1.00 AU for consistency in image capture. Each image was analyzed using a custom Python script (see Supplemental Material) to quantify the number of cells and fluorescence per cell using ImageJ. Post image acquisition, all images were analyzed using Image J. The Python script used details of the general guideline to follow for repeatable measurements for both fluorescence intensity per GFPuv-*E*.*coli* and cell count per image. The code allowed for a repeatable image analysis for an accurate comparison of relative GFP expression per GFPuv-*E*.*coli* and total cell count.

**Fig 2.**
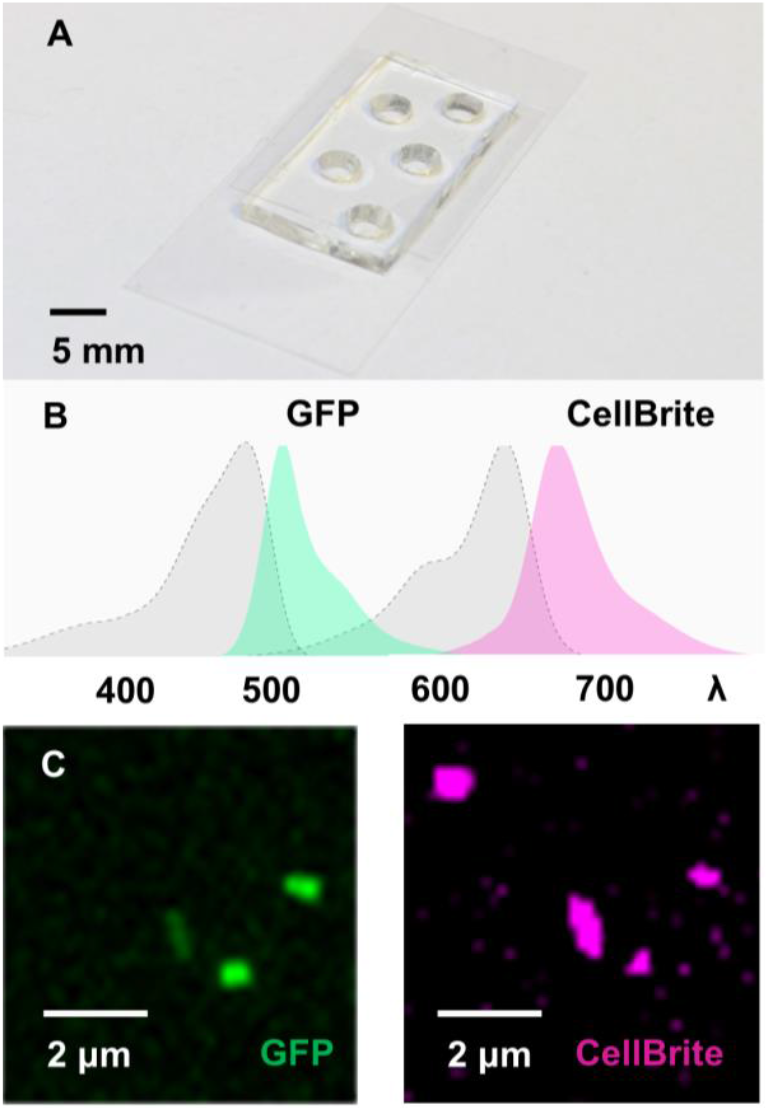
Photograph of microdevice used for confocal imaging of E.coli samples. **A)** The device was manufactured using PDMS bonded to a glass slide. Wells for samples have a diameter of 3 mm. **B)** The peak excitation and emission wavelengths used for this study were λ_ex,GFP_ = 495 nm, λ_em,GFP_ = 509 nm, λ_ex,CellBrite_ = 640 nm, and λ_em,CellBrite_ = 662 nm. **C)** The GFPuv image compared with the CellBrite stained image.

### 2.4 Statistical Analysis

All data points were collected in technical triplicates at various time points over 24 h throughout two independent 300L bioreactor runs. Technical triplicates were compared using a repeated measures ANOVA as well as a Tukey Means Comparison test (α = .05) to determine if there was significant difference between the time point samples. Due to identical bioreactor set up and time points of sample collection, data from both bioreactor runs were averaged and analyzed. This analysis method was used to find the statistically significant highest point for each metric. To assess the strength of correlation between different methods, Pearson’s correlation coefficient was calculated. This analysis compared trendlines across various fluorescence intensity measurement techniques as well as different OD_600_ data collection methods.

## 3. RESULTS & DISCUSSION

### 3.1 Bioreactor Metric Analysis

During the bioreactor run, several inline (pH, DO, Temperature) and at-line (Glucose and OD_600_) parameters were measured throughout the 24 h process, which can be seen in **Figure 3**. The measured variables included optical density (OD), which increased with GFPuv-*E*.*coli* proliferation and glucose which decreased as GFPuv-*E*.*coli* proliferated and consumed. The weight of the bioreactor content was measured using an internal calibrated scale to ensure it was unchanged and maintained within a specific range (275 kg+/-10 kg), although some variability in this measurement existed due to removal of samples throughout the process and standard measuring error. Agitation began in response to changes in DO and the rpm was recorded. The dissolved oxygen content (DO) indicates oxygen available for consumption. Aeration (slpm) is used to promote oxygenation, and agitation (rpm) is introduced to promote both mixing and uniform oxygenation in the culture medium. The pH was measured between 6-7. Temperature and pressure were regulated between 29.8-30.1 ^○^C and 4.95-5.05 psi, respectively. It should be noted that aeration, temperature, and pressure show some variation due to the scale of the presented values; however, there was statistically negligible variation in these parameters. **Figure 3** shows the change in these metrics over time for Run 2.

**Fig 3.**
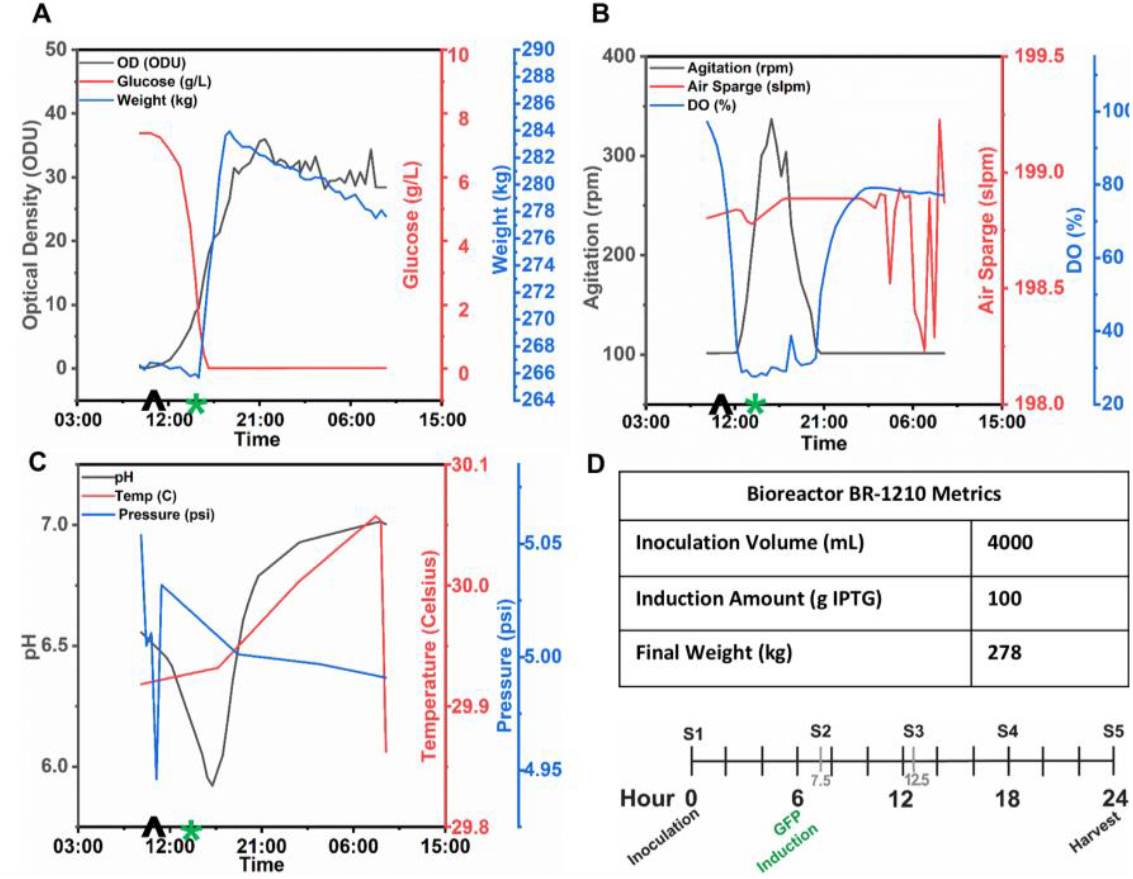
Parameters measured during the bioreactor run, highlighting changes during inoculation at 09:08 (^) and induction at 15:08 (*). **A)** Optical density increased drastically after induction and levelled out afterwards. Glucose levels decreased with growth, as expected. Weight remained within 275 kg +/-10 kg range. Fluctuations in weight are due to standard variability within measurement. **B)** Agitation, Air Sparge and Dissolved Oxygen were all monitored throughout the bioreactor run. The sharp decrease in DO% after inoculation levels out after induction which then increases after an increase in agitation. Air sparge remains at a steady rate and begins to vary towards the end of the process before final harvest. **C)** Changes in pH, temperature and pressure were all tightly monitored and maintained throughout the bioreactor run but do show a decline in pH and pressure after inoculation. Pressure begins to rise and level out immediately after inoculation while pH slowly declines until after induction and then begins to rise and level out until final harvest. The temperature remains within the strict 29.8 and 30.1 Celsius range. **D)** Table 4.D gives a brief overview of other important parameters measured for this run. The timeline in 4.D shows inoculation began at hour 0, GFP induction at hour 6, and final harvest at hour 24. Time points for the five samples analyzed included hour 0, hour 7.5, hour 12.5, hour 18 and hour 24.

Results in **Figure 3A** show a correlation between glucose decline and an increase in optical density and weight (measured by settling of cells on the bottom of the bioreactor). This is expected, as glucose is consumed by cells during the proliferation period. **Figure 3B** depicts changes in dissolved oxygen (DO) as a percentage compared to agitation rpm which is inversely correlated. This relationship can be explained as when agitation and air sparging are introduced to the system, more dissolved air and therefore oxygen is added to the solution. These metrics are important to capture to ensure adequate functionality of the bioreactor and to give insight into the biological processes occurring during inoculation, GFP induction and thereafter.

Glucose levels correlate with cell proliferation because as cells are expanding, they are consuming glucose and therefore levels will decrease as cells proliferate. Optical Density (OD) also relates to *E. coli* proliferation as while cells are proliferating the optical density will increase due to a larger number of cells in the culture. Temperature monitoring is important for ensuring that the environment is suitable for cell proliferation. Other variables such as pressure, pH and DO % must all be tightly monitored and maintained within a certain range for optimal *E. coli* health, protein expression and proliferation. As cells are expanding and consuming glucose and oxygen, the glucose and DO% levels will drop.

### 3.2 Plate Reader Assays

Results from the plate reader for bioreactor runs 1 and 2 are shown in **Figure 4** and are considered offline analysis. OD_600_ and fluorescence intensity of *E*.*coli* suspended in the broth media were measured. Subsequently, the suspension was centrifuged to isolate the supernatant, which was then plated and measured separately. Technical triplicates were measured for each sample. The OD_600_ values relate to cell density while fluorescence intensity relates to GFP expression. The isolation and measurement of the supernatant fluorescence intensity is essential to infer potential GFP leakage from the *E*.*coli* into the broth media. It is well-established that bacterial membrane and cell wall become more permeable as they undergo cellular death, leading to “leakage” in their surrounding media [39]. Therefore, measuring fluorescence intensity in the supernatant can indicate the health of the cell population.

**Fig 4.**
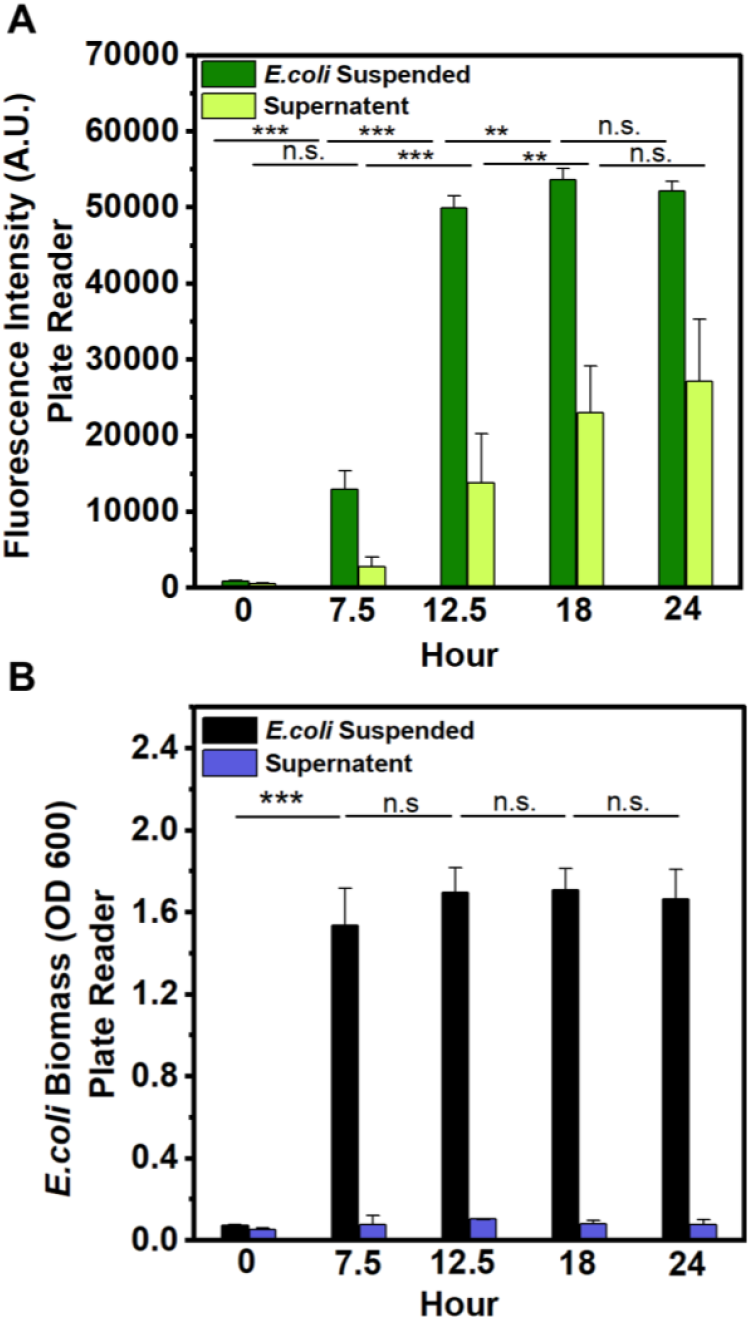
Bar graph representations of fluorescence intensity and OD_600_ for two different bioreactor runs collected via plate reader. **A)** E. coli fluorescence values for 5 samples from bioreactor run 1 and 2 shows a significantly high (p<.001) relative fluorescence intensity of the suspended cell sample at hour 18 which is not significantly different than at hour 24 and a significant (p<.001) increase in fluorescence intensity of the supernatant at hour 18 which is not significantly greater than hour 24. **B)** E. coli OD_600_ values for 5 samples from bioreactor run 1 and 2 shows a significantly high (p<.0001) increase in OD_600_ value of the suspended cell sample at hour 7.5 which is not significantly different than hours 12.5, 18 or 24 indicating that cells proliferated and plateaued in growth after hour 7.5.

The OD_600_ values of the cells suspended in broth, indicative of cell density, show comparable changes in density profiles for both bioreactor runs. There was no significant change in cell density values of the supernatant for both runs, indicating that the supernatant was successfully isolated. The OD_600_ values of the suspended cells indicate the samples taken at hours 12.5 and 18 had the highest (p<.0001) cell density for run 1 and hours 12.5, 18 and 24 had the highest (p<.0001) OD_600_ values for run 2 based on a repeated measures ANOVA and Tukey Means comparison. Run 1 shows no significant difference between OD_600_ values for hours 12.5 and 18 and run 2 shows no significant difference in OD_600_ values between 12.5, 18 and 24 hours. In run 1, there was a significant decline in cell density after hour 18 (p<.0001), suggesting potential cell death occurred. Fluorescence intensity values for bioreactor run 1 indicate a significantly higher (p<.0001) overall fluorescence at hour 18 for suspended cells and hour 24 for supernatant. While GFP expression in bioreactor 2 peaks significantly at hour 12.5 for suspended cells and hour 24 for supernatant.

The OD_600_ and fluorescence intensity data obtained from the plate reader are part of the offline analysis of these bioreactor runs. The observed increase in fluorescence intensity of the supernatant across both bioreactor runs could indicate cell death or an increased membrane permeability, which is the highest at 24 hours (but not significantly higher than hour 18). The fluorescence intensity value of the suspended cells increases and peaks at hour 18 but thereafter does not significantly change at hour 24. These trends underscore the potential benefits of optimizing the duration of future bioreactor runs to enhance total product yield and streamline resource utilization in this bioprocess.

Statistical analysis, based on a Repeated Measures ANOVA with Tukey Means Comparison, was conducted on the technical triplicates from two separate biological runs collected for each time point via the plate reader. Comparison of each time point facilitated the identification of peak values for optical density (correlated with cell density) and fluorescence (associated with GFP expression).

### 3.3 Microscopy and Image Analysis Results

Image analysis was performed as a secondary method for quantification of relative fluorescence intensity, and cell density at each time point. Summary **Figure 5** highlights this offline analysis pipeline from confocal image capture through image analysis for samples collected from both bioreactor runs. Raw images were first collected for every sample using both λ_ex_ = 488 nm; λ_em_ = 507 nm and λ_ex_ = 642 nm; λ_em_ = 662 nm which produced two separate images for each region. Individual cells were identified based on the CellBrite 640 membrane stain. This membrane stain was used as a marker for the *E*.*coli*. Images were collected until a minimum of 200-300 individual GFPuv-*E*.*coli* bacterium were counted in each sample group, where later point samples realized individual counts of ∼1000 for similar image numbers.

**Fig 5.**
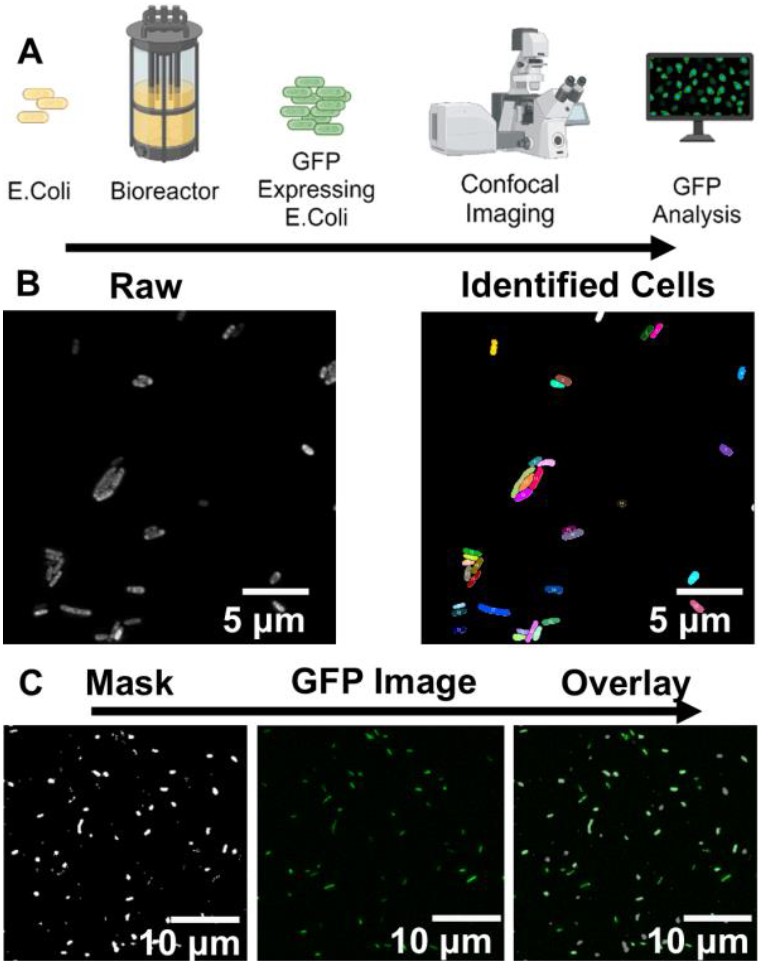
Offline image analytics process for calculating fluorescence intensity per cell and cell count. **A**) Samples are collected from the bioreactor at various time points and analyzed using confocal microscopy. **B)** Raw images collected are processed using Python code in Image J to identify each individual E. coli cell based on the CellBrite 640 membrane stain. **C)** The identified cells then are used as a “mask” which when overlayed with the fluorescence image will measure the relative mean fluorescence intensity per E. coli.

Once individual cells were identified, these identified cells were made into a cell mask. This mask was then overlayed on top of the EGFP image of the same region. Each individual cell, marked with the mask and overlayed on the EGFP image was then measured for fluorescence intensity. The code described earlier allows for repeatable fluorescence intensity measurements for each individual image for all samples. This method for measuring fluorescence allows for a higher resolution offline analysis as compared to the plate reader fluorescence intensity data. The confocal image analysis of fluorescence intensity per cell can be used to measure the distribution of fluorescence per cell as opposed to the overall fluorescence intensity per well measured in the plate reader. Multiple images were collected for each sample to accumulate hundreds of cells for fluorescence intensity analysis.

Image analysis of fluorescence intensity for two separate bioreactor runs can be seen in **Figure 6**. Fluorescent intensity data from both bioreactor runs follows a similar trend. Specifically, the highest mean fluorescence intensity can be found at hour 12.5 (p<.0001). The statistical analysis was performed via repeated measure ANOVA and Tukey Means Comparison with an alpha of .05.

**Fig 6.**
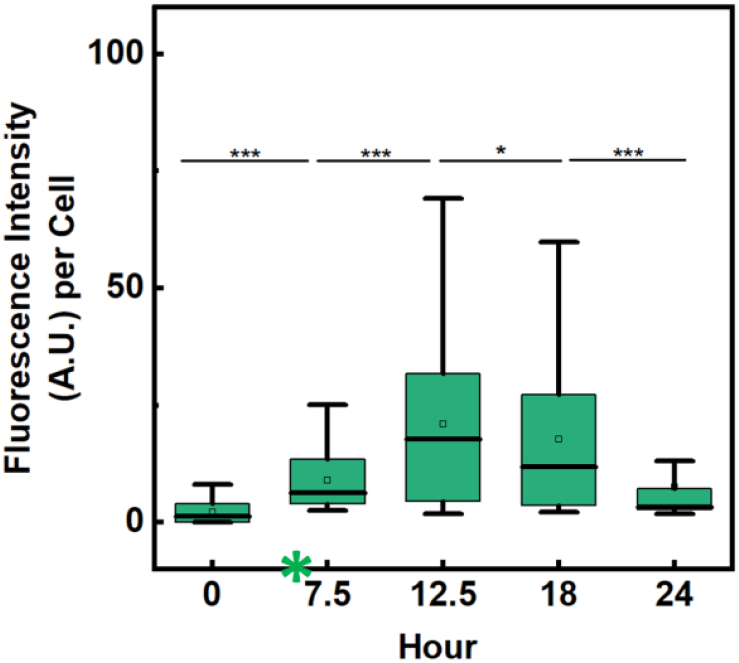
Fluorescence intensity per cell as determined by confocal image analysis for two separate bioreactor runs. The peak of fluorescence intensity (A.U.) per E. coli for both runs is at hour 12.5 (p<.0001) which then begins to significantly decline at hour 18 (p<.05).

Overall, the fluorescence intensity per cell area measured at five different time points (hour 0, 7.5, 12.5, 18, and 24) revealed significant variations (Repeated Measures ANOVA and Tukey Means Comparison, p < 0.0001) between all-time points for the two runs. Fluorescence intensity data shows a distinct peak in fluorescence intensity at the 12.5-hour time point. After fluorescence reached its maximum levels at hour 12.5, the fluorescence intensity begins significantly declining in hours 18 and 24.

These results indicate that peak GFPuv expression per *E*.*coli* is around the 12.5 hour period post inoculation. This peak in fluorescence intensity per cell area is different than the peak in overall fluorescence measured with the plate reader, which showed a peak at hours 18 and 24. This difference can be attributed to the fact that the plate reader takes a singular reading of fluorescence intensity of an entire sample and is much lower resolution than the confocal fluorescence intensity measurement per cell area. Although the overall fluorescence intensity may increase during hours 18 and 24, this does not account for any cellular membrane leakage or levels of proliferation that could impact the final measurement. Overall, the repeatable imaging pipeline and analytics presented in this study are key for multimodal analysis. Additionally, on the single cell analysis level, the imaging technique can be performed on a small number of either fixed or live cells which would allow for retrospective analysis. This imaging pipeline offers an alternative to flow cytometry which is a commonly used PAT for single cell level of analysis [29, 37, 40].

To calculate cell density per sample, a standardized number of images were used. Initially, 56 individual images were captured, and the total cell number was calculated per image using the cell mask process previously described. This cell density quantification method provides a total cell count per image as opposed to an OD_600_ value per well measured in the plate reader. This offline analysis method was used to both give higher resolution and a means of comparison for the data captured through plate reader analysis. Samples from hour 0 were diluted 10-fold less than all other samples due to lack of cell density and therefore cell counts per image were divided by 10.

After images were captured and analyzed, a sample of 50 randomly selected images were compiled to determine the distribution of cell count per image. Additionally, a sample of the remaining 6 images were compiled to determine the distribution of cell count per image. The results of these two sample sizes were compared. The results from the 50-image sample size were more normally distributed than the results from the 6-image sample size and show more significance in the difference between the groups as seen in **Figure 7**.

**Fig 7.**
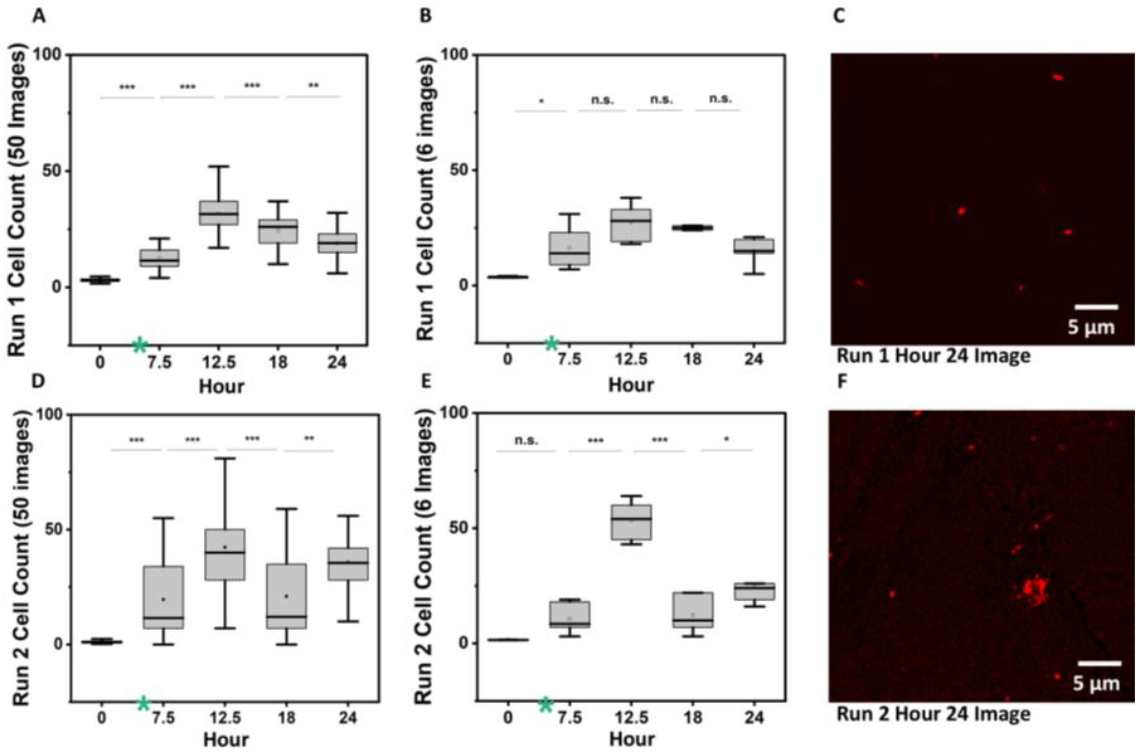
Cell count per image analyzed at different sample sizes for two bioreactor runs. **A)** In run 1, with a 50-image sample size, the peak in cell number per image is observed at hour 12.5 (p<.0001). **B)** In run 1, based on a 6-image sample size, determination of the peak in cell number is inconclusive due to a non-significant difference in cell number when comparing hours 7.5, 12.5, 18, and 24. **C)** Hour 24 image of E.coli from run 1 showing intact E.coli which are fairly sparse. **D)** In run 2, also with a 50-image sample size, the peak in cell number per image is observed at hour 12.5 (p<.0001). **E)** Conversely, in run 2, based on a 6-image sample size, a peak is observed at hour 12.5 for cell number per image (p < .0001). **F)** Hour 24 image of E.coli from run 2 showing cells potentially beginning to die via apoptosis showing multiple cell fragments which can impact the cell mask and cell count.

Results from the 50-image sample size show the highest cell count per image to be at 12.5h for both run 1 (Repeated Measures ANOVA and Tukey Means Comparison, p<.0001) and run 2 (p <.0001). Analysis of 6 image sample size for run 2 shows the highest cell count per image to be at hour 12.5 (p<.0001). However, when analyzing the 6-image sample size for run 1, there is no significant difference between hours 7.5, 12.5, 18 and 24. Therefore, in order to determine the highest cell count per image, more than 6 images are necessary. These results of cell count per image follow a similar trend to the OD_600_ data from the plate reader analysis which both correlate to cell density. This information further supports the claim that there may be a point of diminishing returns during this bioreactor where cell density begins to decrease due to increased cell death. **Figure 7** highlights the cell fragments at hour 24 that begin to appear in Run 2 and can impact the cell mask and subsequent count.

### 3.4 Comparison of Analytical Methods

Single-cell and batch-level analytical methods for the quantification of GFPuv yield were demonstrated and compared. While previous studies have focused on single cell level analysis of GFP expression, utilizing flow cytometry, or have focused on batch-level protein quantification via optical density OD_600_ and fluorescence assays, this study bridges the gap between these two analytical levels [17, 42]. **Figure 8** shows the representative comparative metrics, as illustrated by analysis of Run 2. Analysis of plate reader data, confocal image analysis, and inline bioreactor measurements suggest that the peak *E. coli* density and GFP expression are between hours 12.5-18h. A strong correlation was observed between inline bioreactor and plate reader methods for OD_600_ measurement (Pearson’s correlation coefficient of 0.87). Microscopic image analysis and plate reader methods for quantification of GFPuv yield were also comparable (Pearson’s correlation coefficient of 0.96) for samples collected during the initial 18h of the bioreactor runs. Peak GFPuv expression was expected at 18h.

**Fig 8.**
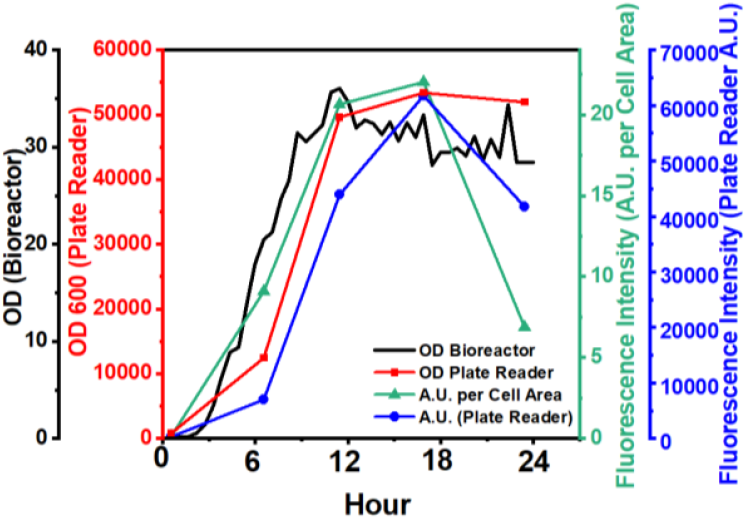
Results from bioreactor Run 2 including both inline and offline quantification of GFPuv yield. OD_600_ data was collected inline from the bioreactor. OD_600_ and bulk fluorescence intensity data were collected using a plate reader, and fluorescence intensity per cell was computed via confocal image analysis. Peak yield was between 12.5-18h.

A surprising outcome was realized after 18h. A steep decline and divergence of outputs from both plate reader assay and microscopic methods was observed, suggesting that as the cells are proliferating, there may be a point where they become overpopulated or underfed and start not only decreasing their rate of proliferation but also begin to die. This idea can be supported by the supernatant fluorescence intensity data, which shows a steady increase in GFPuv expression over time. This increased supernatant fluorescence intensity could indicate the *E. coli* have increased permeability, and GFPuv is leaking into the surrounding media. Apoptosis would also release GFPuv into the media [39, 41]. This effect was most apparent in the microscopy-based method, which when understood in conjunction with when used with batch-level analysis like plate reader measurements, provides crucial additional insights. This analytical pipeline not only identifies key time windows for peak protein expression but also offers insight into cell leakage and viability, as detected through supernatant fluorescence. This data suggests it might be optimal to shorten this bioprocessing timeline after 18h to improve GFP expression and *E*.*coli* viability; however, more data collected from future runs will be necessary to validate this claim. The limitations of this study include sample size, which was constrained by only two biological runs.

The multifaceted analysis and comparison of several metrics gives a higher resolution view of this bioprocess, which can be used for further optimization and enhancement in the future. This short study combines two methods for fluorescence intensity quantification which correlates with GFP expression which could indicate optimal bioreactor settings with further investigation. The microscopy-based method, unlike flow cytometry, requires fewer cells, does not demand specialized equipment or technical expertise, and provides differentiation of debris from target and a rapid inference of lysed cells in the population. [37, 40].

## 4. CONCLUSION

This study underscores the importance of multimodal analytical approaches in bioprocess monitoring to advance Industry 4.0 applications, where scalable and automated process analytics are vital for high-throughput operations. For example, analyzing data at both the batch and cellular levels during the process development stage can identify divergent responses in your process and provide crucial information for accurately setting operational parameters—such as fermenter harvest time—before transitioning to manufacturing. This comparative approach supports more precise process optimization and smoother scale-up to industrial production.

In our study, we highlight this by comparing batch-level analysis with single-cell, semi-automated imaging analysis to measure the GFPuv yield across the entire bioreactor run. This study also offers a rare perspective on pilot-scale production (300L) with step-by-step data collection and analysis methods. A clear point of diminishing returns was identified with offline techniques, while image analysis revealed insights into factors contributing to productivity loss. Batch-level analysis, based on OD_600_ and fluorescence intensity measurements of the supernatant and cell suspension, highlighted protein leakage and changes in overall fluorescence expression. Inline metrics, including OD_600_, pH, and glucose levels, provided real-time insights into the environmental conditions within the bioreactor. All three methods indicate that an earlier harvest could potentially achieve a more efficient product yield.

In conclusion, while inline and at-line metrics offer valuable real-time insights, offline techniques like image analysis provide high-resolution data that are crucial for evaluating cellular health and product distribution—key factors in assessing yield decline, which may signal lysis or cell death. These findings are instrumental for bioprocess optimization, particularly in identifying optimal harvest times, as evidenced by the strong correlations observed between the different analytical methods. While this study was limited by the scheduling constraints of bioreactor runs, future research should incorporate biological triplicates to enhance the robustness and generalizability of the results.

## 5. ACKNOWLEDGEMENTS

G.R was supported by the Chemistry of Life Pre-Doctoral Training Grant at NC State University (NIH Grant 5T32GM141887-03). This work was performed in part at the North Carolina State University BTEC Facility. The authors wish to acknowledge the support provided by the Novo Nordisk Foundation (AIM-Bio Grant NNF19SA0035474), National Science Foundation (Grant EBMS-2033997), the Triangle Universities Center for Advanced Studies Inc. (TUCASI), and the North Carolina Viral Vector Initiative in Research and Learning (NC VVIRAL) at NC State University.

## 6. STATEMENTS AND DECLARATIONS

### Ethical Approval

Neither ethical approval nor informed consent was required for this study.

## Supplementary Information

### Code for Image Analyses

The code used for image analysis is compatible in both Python and Macros languages and should be used with FIJI Image J image analysis software. The description of each steps operation is denoted below with “//.”

open(“.lif”);

*<u>//open up original image which should include both GFP and Cell Mask images</u>*

selectWindow(“A-S1-IM3 - C=0”);

*//Select GFP Image (*DO NOT CHANGE BRIGHTNESS OR CONTRAST)*

run(“Z Project…”, “projection=[Max Intensity]”);

selectWindow(“A-S1-IM3 - C=1”);

*//Select Cell Mask Image (CellBrite 640)*

run(“Z Project…”, “projection=[Max Intensity]”);

run(“Show Info…”);

run(“Brightness/Contrast…”)

*<u>// Adjust brightness and contrast to reduce background noise and enhance cell mask as needed</u>*.

run(“Apply LUT”);

run(“16-bit”);

setOption(“ScaleConversions”, true);

run(“8-bit”);

setAutoThreshold(“Default no-reset”);

run(“Threshold…”);

setThreshold(47, 255);

*<u>// Optimize the threshold to ensure no background noise is included</u>*

setOption(“BlackBackground”, true);

run(“Convert to Mask”);

*<u>// This sets the mask in order to define what is a cell and what is not</u>*

run(“Close”);

run(“Watershed”);

run(“Analyze Particles…”, “size=15-Infinity pixel display exclude clear summarize overlay”);

saveAs(“Results”, “Results.csv”);

selectWindow(“A-S1-IM3 - C=1”);

run(“Set Measurements…”, “area mean min add redirect=[A-S1-IM3 - C=0] decimal=3”);

*<u>// Overlay the cell mask onto the GFP image in order to quantify relative mean GFP per E.coli</u>*

run(“Analyze Particles…”, “size=15-Infinity pixel display exclude clear summarize overlay”);

saveAs(“Results”, “S1Results.csv”);

